# Inferring demographic parameters in bacterial genomic data using Bayesian and hybrid phylogenetic methods

**DOI:** 10.1101/190041

**Authors:** Sebastian Duchene, David A Duchene, Jemma L Geoghegan, Zoe A Dyson, Jane Hawkey, Kathryn E Holt

## Abstract

**Background:** Recent developments in sequencing technologies make it possible to obtain genome sequences from a large number of isolates in a very short time. Bayesian phylogenetic approaches can take advantage of these data by simultaneously inferring the phylogenetic tree, evolutionary timescale, and demographic parameters (such as population growth rates), while naturally integrating uncertainty in all parameters. Despite their desirable properties, Bayesian approaches can be computationally intensive, hindering their use for outbreak investigations involving genome data for a large numbers of pathogen isolates. An alternative to using full Bayesian inference is to use a hybrid approach, where the phylogenetic tree and evolutionary timescale are estimated first using maximum likelihood. Under this hybrid approach, demographic parameters are inferred from estimated trees instead of the sequence data, using maximum likelihood, Bayesian inference, or approximate Bayesian computation. This can vastly reduce the computational burden, but has the disadvantage of ignoring the uncertainty in the phylogenetic tree and evolutionary timescale.

**Results:** We compared the performance of a fully Bayesian and a hybrid method by analysing six whole-genome SNP data sets from a range of bacteria and simulations. The estimates from the two methods were very similar, suggesting that the hybrid method is a valid alternative for very large datasets. However, we also found that congruence between these methods is contingent on the presence of strong temporal structure in the data (i.e. clocklike behaviour), which is typically verified using a date-randomisation test in a Bayesian framework. To reduce the computational burden of this Bayesian test we implemented a date-randomisation test using a rapid maximum likelihood method, which has similar performance to its Bayesian counterpart.

**Conclusions:** Hybrid approaches can produce reliable inferences of evolutionary timescales and phylodynamic parameters in a fraction of the time required for fully Bayesian analyses. As such, they are a valuable alternative in outbreak studies involving a large number of isolates.

## Background

Genomic data are increasingly used to investigate infectious disease outbreaks caused by microbial pathogens. Recent developments in sequencing technologies have made it possible to obtain data for a very large number of samples, at low cost and within a very short timeframe. Phylogenetic methods can make use of these data to infer their evolutionary dynamics, known as phylodynamic inference. For example, genome data obtained during the first months of the 2013-2016 Ebola virus epidemic were used to determine the time of origin of the outbreak and the basic reproductive number (*R_0_*) of the circulating strains [1,2]. Some of the key requirements for these inferences are that the data must have sufficient genetic diversity and that they should be a representative sample of the circulating strains.

Serially sampled data are particularly useful because their sampling times can be used to calibrate the molecular clock. This consists of calculating the rate of evolution, which is the amount of genetic change that has accumulated per unit of time. The rate of evolution is key to infer an evolutionary timescale, typically represented by a phylogenetic tree where the branch lengths correspond to time, known as a chronogram. Some methods assume that the rate of evolution is constant over time, known as a strict molecular clock, but popular Bayesian implementations, such as that in BEAST [3,4], include relaxed-clock models that use a statistical distribution to describe rate variation across time and lineages (reviewed in [5]). Phylodynamic models can be used to estimate the epidemic growth rate (*r*), *R_0_*, and other parameters [6,7]. Importantly, these models describe the expectation of the distribution of node times in the chronogram. As such, inferences drawn from phylodynamic models rely on accurate estimates of evolutionary rates and timescales. A number of statistical methods are available to assess the robustness of inferences of evolutionary rates and timescales; those that are most widely used are implemented under a Bayesian framework (reviewed in [8]).

Bayesian phylogenetic approaches allow sophisticated evolutionary models to be specified. For example, the evolution of a pathogen during an outbreak can be defined as an exponentially growing population with considerable evolutionary rate variation among lineages; which can be modelled by specifying a nucleotide substitution model, a relaxed-clock model and an exponential-growth tree prior. The parameters for all these models are obtained simultaneously and their estimates correspond to posterior probability distributions, such that their uncertainty is a natural by-product of the analysis. Bayesian methods require specifying a prior distribution for all parameters. Although specifying a prior distribution is not trivial for some parameters, their influence can be assessed by comparing them to the posterior. An advantage of specifying prior distributions is that it is possible to include previous knowledge about the data. As a case in point, a known probability of sampling can be represented with a prior distribution in birth-death models [9].

Whilst Bayesian phylogenetic methods have many desirable properties, analysing large genomic data sets under complex models is often computationally prohibitive (e.g. [10,11]). An alternative to full Bayesian methods is to conduct the analysis in several steps. In this hybrid approach the phylogenetic tree, evolutionary rates and timescales, and demographic parameters are estimated separately.

Phylogenetic trees can be rapidly estimated using various maximum likelihood implementations [12–15]. These methods assume a substitution model, but not a molecular-clock or demographic model, such that the branch lengths of the trees represent the expected number of substitutions per site, and are known as phylograms.

Next, phylograms can be used to estimate evolutionary rates and chronograms, for example, using a recently developed molecular clock method based on least-squares optimisation, called LSD (Least Squares Dating) [16]. LSD is more computationally tractable than Bayesian molecular-clock methods, such that it is feasible to analyse genomic data sets with thousands of samples. Although LSD assumes a strict molecular clock, its accuracy is frequently similar to that obtained using more sophisticated Bayesian clock models [17]. Other non-Bayesian molecular-clock methods have also been developed recently with the purpose of analysing large genomic data sets [18–20].

Finally, a range of tools are available to infer phylodynamic parameters from a chronogram, such as that obtained using LSD. For example: TreePar uses maximum likelihood to fit birth-death and skyline models [21]; BEAST2 [4] and RevBayes [22] can fit a range of birth-death, coalescent, and Skyline models using Bayesian inference [7]; and approximate Bayesian computation (ABC) approaches that use tree summary statistics have recently been developed to fit phylogenetic epidemiological models [23,24]. The main disadvantage of these approaches over those that are fully Bayesian is that the estimates are based on a single tree, such that uncertainties in tree topology, branch lengths, and evolutionary rates are ignored. A potential solution is to repeat the analysis using non-para metric bootstrap replicates, but combining the different sources of uncertainty under this framework is not trivial.

Here, we compare the following two methods to infer evolutionary rates and timescales, and demographic parameters:

i. The fully Bayesian method, implemented in BEAST2, to simultaneously infer the phylogenetic tree, evolutionary timescales and phylodynamic parameters;
ii. The hybrid method: phylogram inference using maximum likelihood in PhyML V3.1 [14], chronogram inference using LSD v0.3, and estimation of phylodynamics parameters in BEAST2 using Bayesian inference.

To compare the performance of these two methods, we analysed previously published whole genome SNP bacterial data sets of *Mycobacterium tuberculosis* Lineage 2 [25], *Vibrio cholerae* [26], *Shigella dysenteriae* type 1 [11], and *Staphylococcus aureus* ST239 [27]. Because these data sets have small numbers of samples (n=63 for *M. tuberculosis*, n=122 for *V. cholerae*, n=121 for *S. dysenteriae*, and n=74 for *S. aureus*) their analyses are computationally tractable using both approaches. We also demonstrate the unique potential of the hybrid approach by analysing two genomic data sets with larger numbers of sequences, which have been difficult to analyse using a fully Bayesian approach; a global sample of *S. dysenteriae* type 1 (n=329) and *S. dysenteriae* type 1 lineage IV (n = 208) [11]. Finally, we validated the performance of the hybrid approach using a simulation experiment.

## Results

### Estimates of evolutionary rates and timescales

We compared estimates of rates and evolutionary timescales using the full Bayesian approach in BEAST2 and LSD. Because our data consist of SNPs, we used ascertainment bias correction by specifying the number of constant sites from the core genome. In BEAST2 we used both the strict and the uncorrelated lognormal (UCLN [28]) clock models. We investigated the degree of rate variation among lineages by inspecting the coefficient of rate variation, estimated in the UCLN model. This parameter is the standard deviation of branch rates divided by the mean rate. The data are considered to display clocklike behaviour if the distribution for this parameter abuts zero. Therefore, we used this parameter to select the clock model in BEAST2 for each data set, as suggested in previous studies [29,30]. The *M. tuberculosis* data set was the only data set to support a strict clock over the UCLN model, whereas the remaining data sets favoured the UCLN model (Fig.1). We set uniform prior distributions for the clock rate, the growth rate (*r*)and the scaled population size (*Φ*). In the context of pathogen evolution, rdetermines the speed of spread of the pathogen in the host population, while *Φ* is proportional to the infected host population size at present.

**Figure 1.**
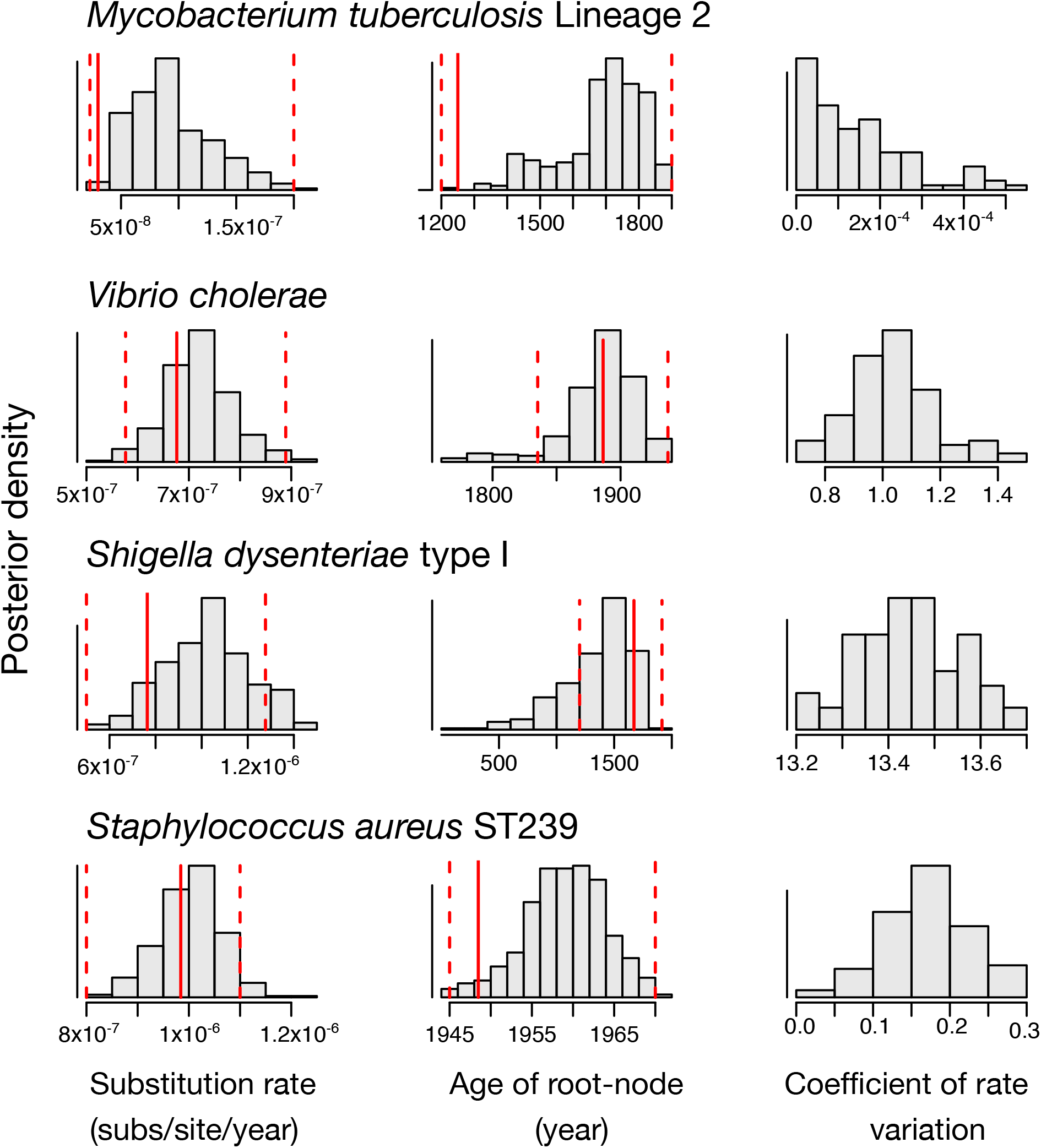
Estimates of evolutionary rate, time to the most recent common ancestor, and the coefficient of rate variation of the UCLN. The histograms correspond to the posterior distribution in BEAST2 using the full Bayesian approach. With the exception of the *Mycobacterium tuberculosis* data set, we used the UCLN clock model because the coefficient of rate variation was not abutting zero. The red solid line is the estimate from LSD, and the dashed lines correspond to the 95% confidence interval. Note that the coefficient of rate variation is not computed for LSD, which assumes a strict molecular clock.

The estimates of evolutionary rates and timescales from these different methods were largely congruent (Fig.1). In all four cases, the 95% credible intervals for the evolutionary rate and age of the root node obtained with BEAST2 overlapped with the 95% confidence intervals obtained for the same parameters with LSD (Fig.1). However, we observed some differences in the mean evolutionary rate estimates, with the estimates from BEAST2 consistently producing higher values than those from LSD. The largest difference in mean rate estimates was observed in *M. tuberculosis*, with a mean rate of 9.37×10^−8^ (95% credible interval: 4.25×10^−8^−1.73×10^−7^) using BEAST2, and 1.10×10^−8^ (95% confidence interval: 1.00×10^−10^−2.02×10^−7^) in LSD (see Fig.1). In contrast we found more congruent mean rate estimates in the *V. cholerae* data set, with estimates of 7.20×10^−7^ (95% credible interval: 5.87×10^−7^−8.65×10^−7^) for the BEAST2 and 6.76×10^−7^ (95% confidence interval: 5.76×10^−7^−8.89×10^−7^) for LSD. The differences in estimates of the root-node age were similar, with the largest difference in the mean root-node age found in *S. aureus* ST239 (mean root-node age of 1958 for BEAST2 and 1949 for LSD) (Fig.1). In most cases, the estimates from BEAST2 were more uncertain with credible intervals that were wider than the confidence intervals from LSD. We investigated two aspects of phylogenetic data that can affect estimates of evolutionary rates; the topological uncertainty and the degree of clocklike variation. We found that the maximum likelihood trees were highly supported, according to local likelihood ratio tests (aLRT) [31] (which ranges from 0 to 1, for low to high branch support, respectively). The median aLRT values across nodes were 0.9 for *M. tuberculosis*, 0.83 for *V. cholerae*, 0.99 for *S. dysenteriae* type 1, and 0.92 for *S. aureus*.

### Assessing temporal structure using a date-randomisation test

We assessed the reliability of our estimates of evolutionary rate and timescales by conducting a date-randomisation test [32,33]. The motivation of this test is similar to that of root-to-tip regressions implemented in TempEst [34]. That is, to determine whether there is sufficient sampling in the data. However, root-to-tip regressions should be interpreted for visual inspection, as opposed to date-randomisations, which are a formal statistical test. The date randomisation test consists in repeating the analysis several times after randomising the sampling dates. The resulting rate estimates correspond to the expected values if there is no association between sampling times and genetic divergence. The data are considered to have strong temporal structure if the rate estimate obtained using the correct sampling times is not contained within the range of values from the randomisations. In a Bayesian context, 10 to 20 randomisations appear to be sufficient [33,35]. We conducted this test in BEAST2 using 20 randomisations and in LSD using 100 randomisations (Fig.2). Interestingly, the results from both tests were congruent, and consistent with visualisations of clock-like behaviour of the data using root-to-tip regressions (Fig.S1). The *M. tuberculosis* data set had no temporal structure with either method (Fig.2): the credible interval of the Bayesian estimate with the correct sampling times overlapped with those from all of the randomisations; using LSD, the estimate with the correct sampling times was around the lower threshold in the program, at 1.00×10^−10^ subs/site/year, which also corresponds to the value obtained for most of the randomisations. The other data sets showed strong temporal structure with both date-randomisation tests: the Bayesian credible intervals using the correct sampling times did not overlap with those from any of the randomisations, and the estimates from LSD using the correct sampling times were not contained within the distributions of the 100 randomisations (Fig. 2).

**Figure 2.**
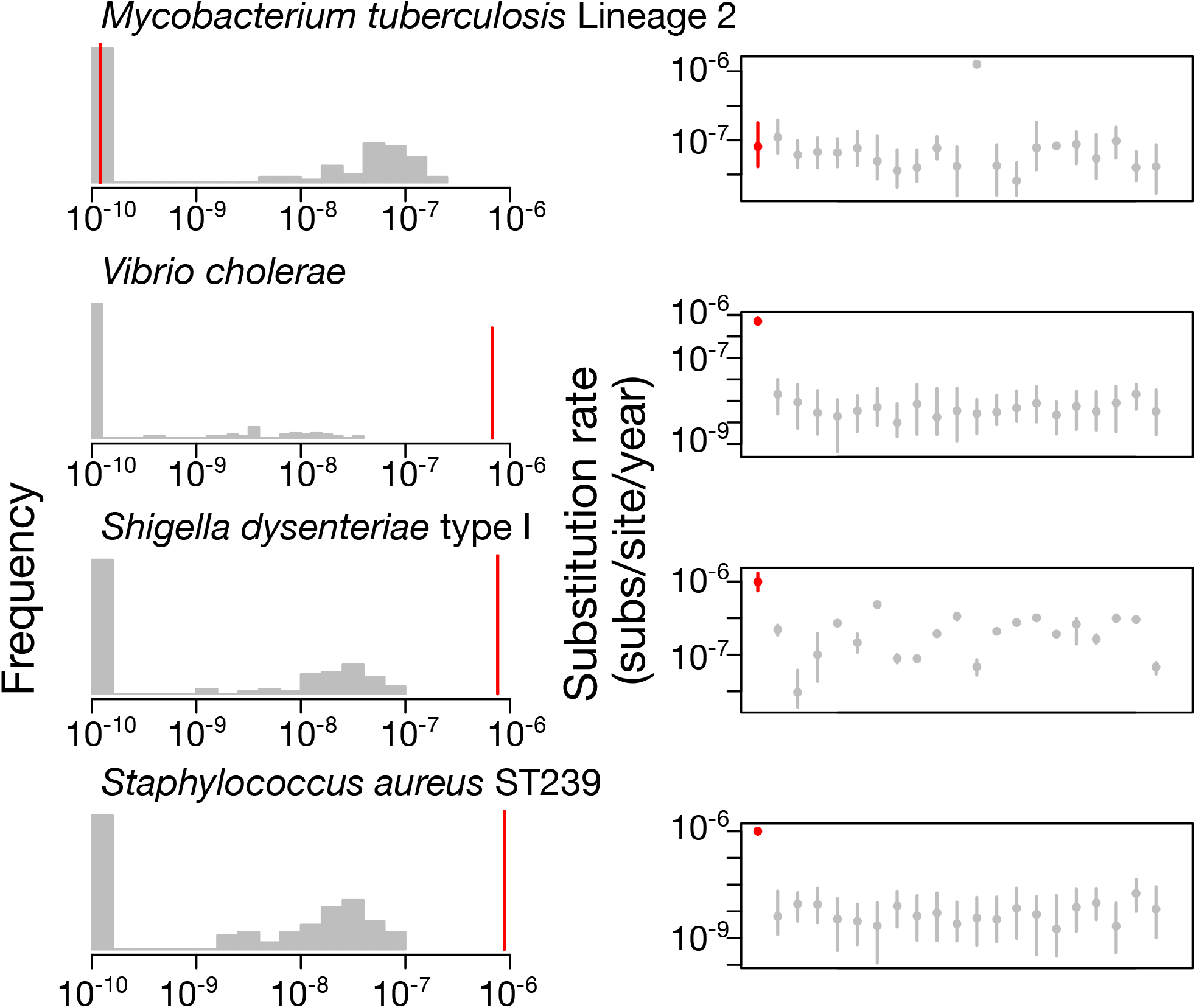
Date randomisation test using LSD and BEAST2. The left column shows histograms of the rate estimates with randomised sampling times in LSD (grey). The red line corresponds to the estimate using the correct sampling times. The right column shows the date randomisation test in BEAST2. The grey bars denote the 95% credible intervals of substitution rate estimates from the randomisations. The red lines correspond to the 95% credible interval of the rate estimates using the correct sampling times. The circles denote the mean value. The *x*-axis in the left column and the *y*-axis in the right column are in logarithmic scale.

### Inference of phylodynamic parameters

We analysed the data sets using the exponential-growth coalescent model in BEAST2, which has two parameters, *r* and *Φ*. Because these are compound parameters, they cannot be interpreted in an absolute scale without additional information about the size of the infected host population at present [36]. In most cases, the posterior distributions of both parameters were very similar when using either BEAST2 or the hybrid approach, with similar means and uncertainties (Fig.3). Although the intervals overlapped in *V. cholerae*, *S. dysenteriae*, and *S. aureus*, the mode of the posterior distribution of *Φ* was higher when using the hybrid approach. The posterior distributions of *r* were almost identical across methods for the three data sets with temporal signal (Fig.3). The uncertainty in estimates of this parameter did not include 0, except in the case of *V. cholerae*, suggesting that most of these bacterial data sets were undergoing population growth. Interestingly, the *M. tuberculosis* data set, which had no temporal structure, was the only data set to display large differences in estimates among the methods (Fig.3).

**Figure 3.**
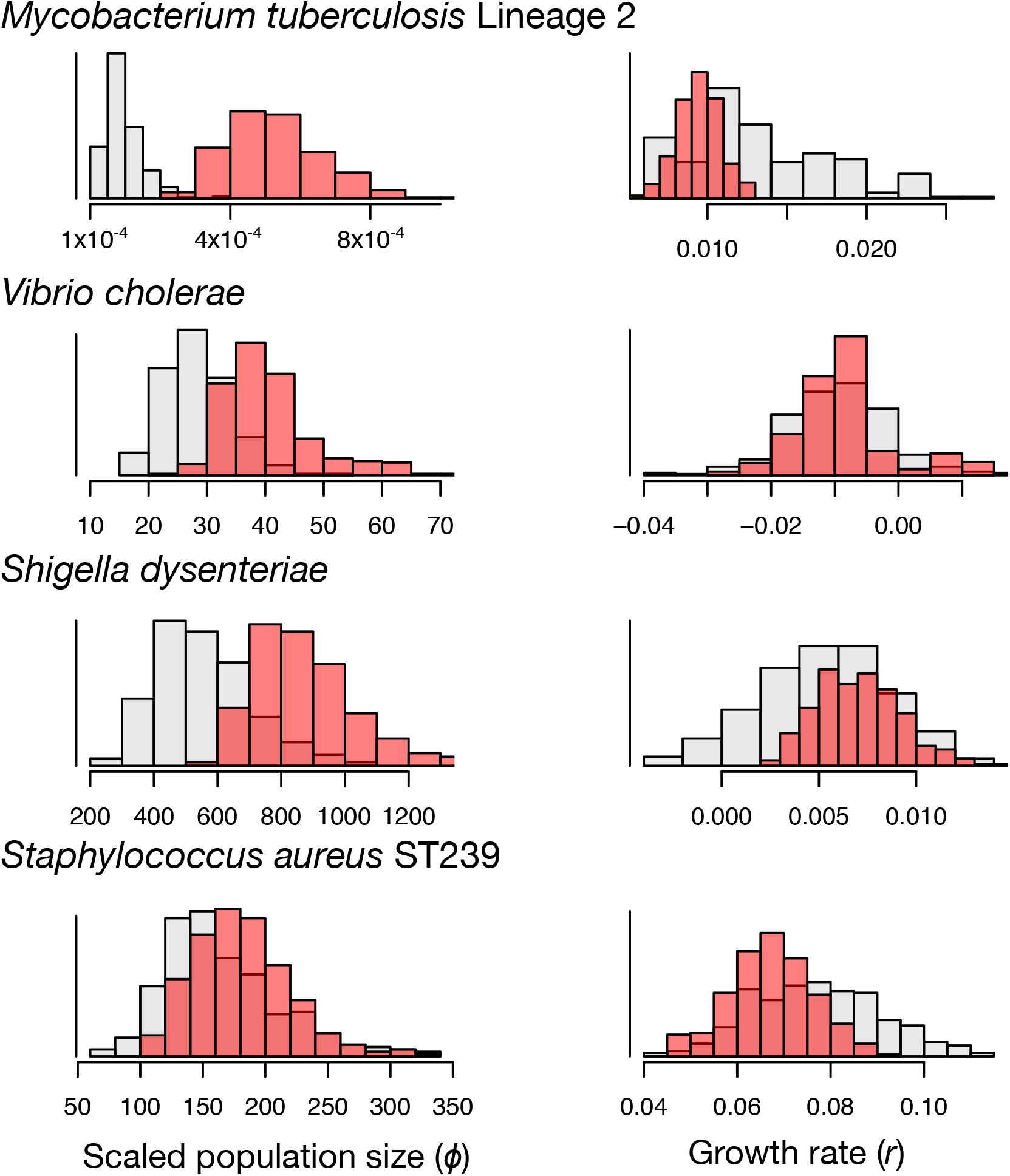
Posterior estimates of demographic parameters, *Φ and r* using the full Bayesian and hybrid approaches. The red histograms correspond to the estimates from the hybrid approach, where the coalescent likelihood is calculated on a fixed tree. The grey histograms correspond to the posterior estimates using the full Bayesian method.

### Application: analysing large data sets using the hybrid approach

Having demonstrated good performance of the hybrid approach on small data sets with strong temporal signal, we applied it to analyse two published genome-wide SNP data sets whose sample size was prohibitively large to analyse under a full Bayesian framework in the original publication. These data sets consisted of: (i)329 samples of *S. dysenteriae* type 1 from [11], which included BEAST2 analysis of a subset of 125 samples; and (ii) 208 samples of lineage IV of *S. dysenteriae* type 1, which was represented by 61 samples in the BEAST2 analysis in the same study [11]. These three data sets displayed strong temporal structure according to the date-randomisation test in LSD, with rate estimates that were not contained within the range of estimates from 100 date-randomisations (Fig.4). The evolutionary rate estimates from LSD were 5.93×10^-7^(95% confidence interval: 3.65×10^−7^−1.65×10^−6^) subs/site/year for *S. dysenteriae* type 1, and 7.04×10^−7^ (95% confidence interval: 3.92×10^−7^−1.54×10^−6^) subs/site/year for *S. dysenteriae* type 1 Lineage IV (Fig.4). Interestingly, the estimate of *r* for *S. dysenteriae* type 1 lineage IV was over an order of magnitude higher than that for the global data set of this bacterium, with a mean of 2.00×10^−2^ for lineage IV compared with 3.40×10^−3^ for the global data set. Importantly, the posterior distributions of *r* for these three data sets did not include zero, indicating epidemic growth (Fig.4).

**Figure 4.**
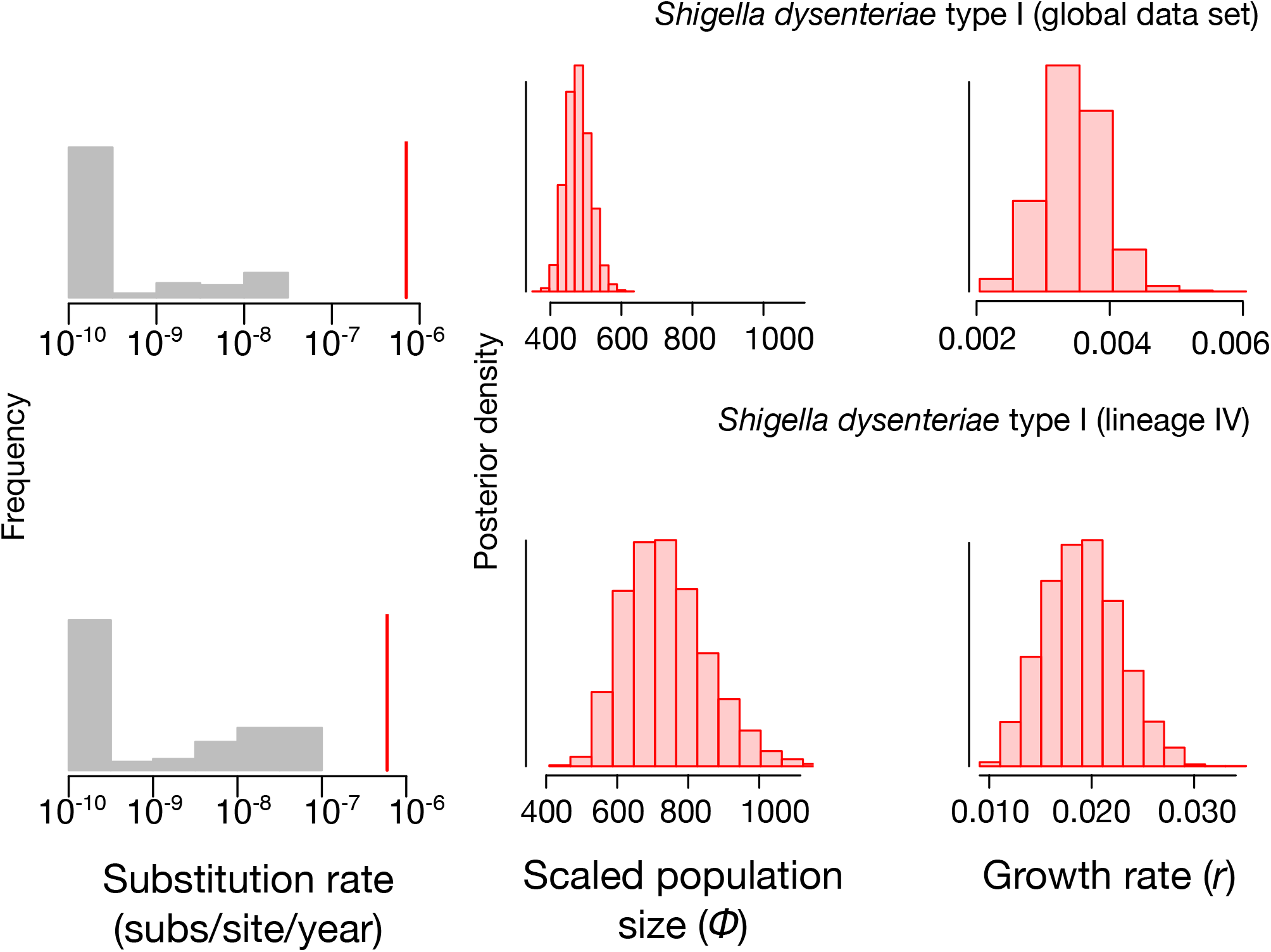
Date randomisation test in LSD and estimates of demographic parameters for large data sets using the hybrid approach. The grey histograms correspond to rate estimates from the randomisations, while the red lines correspond to the estimates using the correct sampling times. The red histograms correspond to the posterior distribution of parameters *Φ* and *r*.

### Validation using simulations

Although our empirical analyses suggest that the hybrid and the full Bayesian method can produce largely congruent results, it is unclear whether the methods are accurate. That is, whether they can recover the true parameter estimates. To investigate this, we conducted a simulation experiment. We simulated 100 whole genome data sets using similar parameters to those we inferred for our *S. dysenteriae* data set. We extracted the SNPs from the synthetic genomes and analysed them using the hybrid and full Bayesian methods, with the same settings that we used for the empirical data. Our date-randomisations in LSD indicated that all of these data sets had temporal structure, with p-values of 0.00. The estimates for the age of the root-node from both methods were very similar. However, it is important to note that our hybrid method uses a single tree, such that the age of the root-node is a point value, whereas the full Bayesian analyses include uncertainty in this parameter. Accordingly, the estimates from LSD were very close to those used to generate the data (within 5 years of the true value), and those from the full Bayesian method always included the true value within their credible interval. The estimates for the demographic parameters, *r* and *Φ*, had credible intervals that always included the true value for both methods, with mean values that often matched those used to generate the data (Fig. 5a). Interestingly, in 10 randomly selected simulation replicates, we found that the credible intervals for the demographic parameters were very similar for both methods, with the hybrid approach sometimes producing more precise estimates. We found no estimation biases in any of the methods (Fig. 5a).

**Figure 5.**
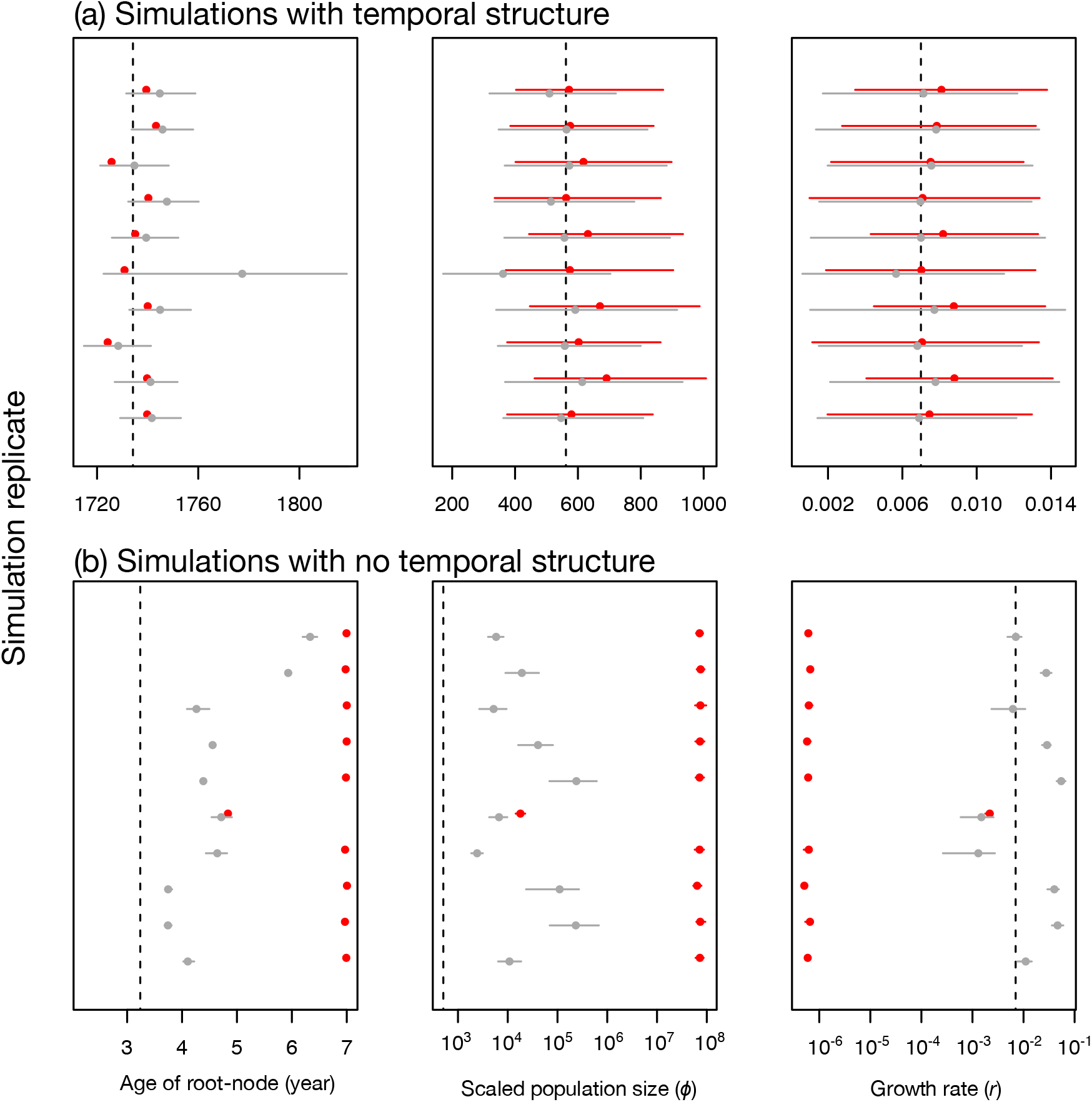
Parameter estimates for 10 randomly selected simulations (from a total of 100). Simulations with strong temporal structure (a) had a *p*-value for the date randomisations test of 0.00, where as those with no temporal structure (b) had a *p*-value of 1. Each row within each panel is for a simulated genome analysis. Estimates in red were obtained using the hybrid method, while those in grey are for the full Bayesian approach. The circles correspond to the mean value, except for the age of the root-node for the hybrid approach (LSD), where it is the point estimate. The bars denote the 95% credible interval. The dashed lines are the value used to generate the data. Note that the *x*-axes in (b) are in log_10_ scale.

We conducted a second set of simulations of data with no temporal structure. To do this, we generated similar sequence alignments as described above, but we assigned random sampling times in our analyses in LSD and in BEAST2. This means that the molecular clock calibration is effectively uninformative. The age of the root-node was overestimated by both methods. In LSD this bias was of over three orders of magnitude, whereas in BEAST2 it ranged between half and three orders of magnitude. The value of *Φ* was similarly overestimated in both methods. The growth rate, *r*, was underestimated by several orders of magnitude with the hybrid approach, but it tended to be overestimated with the full Bayesian method (Fig. 5b). A key result about the simulations with no temporal structure is that *Φ* was always incorrectly estimated, and the true value of *r* was only contained within the 95% credible interval in about 14% of the analyses using the full Bayesian method. Moreover, the estimates with the hybrid approach often displayed larger discrepancies with the correct values.

### Computational demands of the Bayesian and the hybrid methods

The hybrid approach was several times faster than the full Bayesian approach. For example, the computation time for each randomisation of the *V. cholerae* data set each was about 2 hours using BEAST2, where as those in LSD took 1.23 seconds (sec). However, a key aspect of the date-randomisation test in LSD is that the tree topology and branch lengths are fixed for all randomisations, where as they are re-estimated for each randomisation in BEAST2. For the *V. cholerae* data set, a complete analysis using the hybrid approach took: 10.06 minutes (min) to infer a maximum likelihood tree in PhyML, 1.23 sec to estimate the evolutionary rate and timescale in LSD, and 5 min to infer *r* and *Φ* in BEAST2 to obtain effective sample sizes (ESS) of over 200 for all parameters (drawing 1×10^7^ steps, with 1 minutes per 10^6^ steps), for a total of about 15 min, and 1/12^th^ of the time required in BEAST2. Analysis of the full *S. dysenteriae* dataset from [11], the largest data set in our study, took 10.6 sec to analyse in LSD and 1 hour infer *r* and *Φ* BEAST2 (drawing 5×10^7^ steps, with 1.2 minutes per 10^6^ steps), for the 329 sampled sequences.

## Discussion

Our results demonstrate that, as long as a strong temporal signal is present, the hybrid and fully Bayesian methods can produce congruent estimates of evolutionary parameters, even in cases where the data display substantial rate variation among lineages. These methods also yielded similar estimates of demographic parameters in data sets with strong temporal signal, indicating the hybrid approach is a reliable alternative to full Bayesian analyses. However, *r* appears to be more robust than *Φ* to mild differences in estimates of the rate and timescale. This probably occurs because the age of the root-node plays an important role in the population size under the coalescent. In particular, the effective population size, and therefore *Φ*, are known to scale positively with the age of the root-node [37].

Obtaining congruent estimates between the two methods depends on whether the data meet certain criteria. In practice, it is important to verify that the trees have high branch support and that the data have strong temporal structure. The trees inferred here were highly supported, but it is likely that the hybrid approach will produce misleadingly precise estimates (i.e. with narrow confidence intervals) if branch support is low, because the demographic parameters will still be conditioned on a single, and possibly incorrect, tree obtained in step 1 that does not capture uncertainty in the topology. In contrast, in such circumstances the Bayesian method will simply integrate over phylogenetic uncertainty and yield wider credible intervals. Our simulations illustrate ideal conditions, in which the data evolve under the correct model and have strong temporal structure. In this case, we find that both methods produce accurate estimates with similar precision.

Our simulations of data with no temporal structure demonstrate, not only that the hybrid and full Bayesian methods will produce different estimates, but that they both tend to be inaccurate. In the absence of temporal structure, LSD often produces rate estimates at the lower threshold of the program, which was 10^−10^ here. This means that the timescale of the chronogram is overestimated. The value of *Φ* is also overestimated, which occurs because this parameter scales positively with the age of the root-node [37]. Although, we found that *r* was also overestimated, this parameter is determined by the distribution of branches in the tree, such that its error is less predictable. The full Bayesian method produced estimates with smaller bias. We used uniform priors for *Φ* and *r*, and the prior for the age of the root was determined by the coalescent prior. It is likely that these parameters, especially *Φ*, will be affected by different choice of priors. For empirical data with low temporal structure, the hybrid approach will likely be misleading because it is conditioned on a single tree which is probably incorrect. In such cases, it may be necessary to use the full Bayesian method approach because it is possible to include sources of molecular clock calibration via prior parametric distributions, at the expense of much higher computational demands. For instance, a reasonable calibration on the age of the root-node might be sufficient to overcome low temporal structure and to obtain reliable estimates for *Φ* and *r*. To investigate this, it is important to verify that there exists a difference between the prior and posterior for parameters of interest (see Boskova et al. [38] for an investigation of the prior and posterior in Bayesian phylodynamics).

Our results show that the date-randomisation test in LSD appears to be as effective as it is in BEAST2, with the advantage of being much less computationally demanding. As a result, it is possible to use a larger number of replicates, which can improve the power of the test. Moreover, the sampling times under a Bayesian analysis of sequentially sampled data are informative about the tree topology. That is, they impose a high prior probability on trees that cluster sequences with similar sampling times, which can render the date-randomisation test unreliable, with an increase in type I error [39]. Moreover, in some phylodynamic models, the estimate of the age of the root-node and the evolutionary rate are determined by a combination of the sequence data and their sampling times [38], such that assessing temporal structure via the date randomisation test is not trivial. The date-randomisation test in LSD does not suffer from these problems because sequence data alone, not tip dates, are used to infer the tree topology in maximum likelihood.

Critically, the rates estimated using the date-randomisation in test in LSD are not necessarily unimodal in their distribution. This occurs because a lack of temporal structure usually leads to very low rate estimates, which affects randomisations in LSD and in BEAST2. In the case of LSD, very low values for the rate will correspond to the lower threshold set in the program [17], which we arbitrarily set at 10’“subs/site/year, such that most randomisations will have this value. As such, a reasonable approach to interpret the date-randomisation test in LSD is to ensure that the rate estimate with the correct sampling times is higher than those from at least 95% of the randomisations, following the frequentist one-tailed *p*-value of α=0.05.

## Conclusions

As shown here, hybrid methods offer an attractive alternative to full Bayesian approaches for genome-scale data sets with very large numbers of samples. The accuracy and precision of both methods are comparable, but hybrid methods can perform an analysis in a about an eighth of the time required for full Bayesian analyses. Nevertheless, some steps of the hybrid method used here require oversimplifications of the evolutionary process. For example, LSD always assumes a strict molecular clock, such that it is impossible to assess among-lineage rate variation or to pinpoint potential biological causes for why lineages have different rates. The choice of whether to use a hybrid method should be made based on what parameters a user wishes to interrogate. In the context of molecular epidemiology, demographic parameters (*r* and *Φ*) and divergence time information are of primary interest, all of which appear robust to some among-lineage rate variation.

In this study, we used a simple demographic model, the exponential-growth coalescent. This model appears to be well suited when outbreak data are sampled at an early stage, but it makes several assumptions, including that the population of susceptible hosts is constant and that there is no population structure [6], A better understanding of the data used here requires more sophisticated phylodynamic models, such as those that include changes in diversification parameters over time [40],and migration [41]. To this end, our results suggest that harnessing the power of such models and large-scale genome sequencing can be done through hybrid approaches.

## Materials and Methods

### Data collection

Our bacterial data sets consisted of publically available genome data. We obtained all of our genome-wide SNP alignments from a previous studies [11,25,27,35]. These data sets are freely available online (github.com/sebastianduchene/bacteria_genomic_rates_data). These data have had regions with evidence of recombination removed using Gubbins v2 [42].

### Phylogenetic analyses under the fully Bayesian approach

We analysed the sequence alignments in BEAST V2.4 using the sampling times for calibration, the GTR+Γ substitution model, the exponential-growth coalescent tree prior, and two clock models; the strict and the UCLN. We used the default priors for all parameters. Our Markov chain Monte Carlo (MCMC) sampling scheme consisted of a chain length of 5×10^8^ steps, sampling every 10^4^ steps. We verified that the ESS for all parameters was at least 200. To determine whether the data had temporal structure, we conducted a date-randomisation test by randomising the sampling dates 20 times and repeating the analyses [33].

### Phylogenetic analyses using the hybrid approach

We inferred phylogenetic trees using maximum likelihood in PhyML v3.1. We used the GTR+Γ substitution model, and a search strategy that combines the nearest-neighbour interchange and subtree prune and regraft algorithms. To assess branch support, we calculated the aLRT score for each branch. To visually assess temporal structure, we conducted a regression of the root-to-tip distances as a function of the sampling times using TempEst v1.5 [34]. To determine the optimal root in this program we selected the position that maximised R^2^.

We analysed the maximum likelihood trees (i.e. phylograms) in LSD v0.3 to infer the evolutionary rate and timescale. We set the sampling times as calibrations and allowed the program to determine the optimal position of the root. We constrained the branching times of the estimated chronograms such that daughter nodes must be younger than their parent nodes. To obtain an uncertainty around estimates of times and rates, we conducted 100 parametric bootstrap replicates of the branch lengths, as implemented in the program. Therefore, the uncertainty corresponds to the 95% confidence interval of the parametric bootstrap values. We conducted a date-randomisation test 100 times by randomising the sampling times in the ‘date’ file and running LSD each time. In this version of the test, the phylogenetic tree topology and branch lengths are fixed.

We used the chronograms estimated in LSD to infer demographic parameters in BEAST2. This consists in setting the input file to calculate the posterior as the likelihood of the tree given the model parameters multiplied by the priors on the parameters. In the exponential growth coalescent there are two parameters; *Φ* and *r*. We used an MCMC chain length of 1×10^7^ sampling every 10^4^ steps, and we verified that all parameters had ESS values of at least 200.

### Simulations

We simulated whole genome sequence alignments using the parameters from our *S. dysenteriae* data set. To do this, we took the highest clade credibility tree from this data set inferred in BEAST2 and simulated the evolutionary rate using NELSI [29], according to an UCLN clock model. We used a mean rate of 10^−6^ subs/site/year and a standard deviation of 10^−7^. We used Seq-Gen v1.3 [43] to simulate genome sequence alignments of 3,750,125 nucleotides using the GTR+Γ substitution model with the mean parameter estimates for the empirical *S. dysenteriae* data. Finally, we extracted the SNPs from these alignments and analysed using the same method as for our empirical data. For our simulations with no temporal structure we set random sampling times for our analyses in LSD and BEAST2. In all cases, we conducted a date-randomisation test in LSD, as used in our empirical data analysis.

## Abbreviations

LSD: Least-squares dating
ABC: Approximate Bayesian Computation
UCLN: uncorrelated lognormal clock
aLRT: local Likelihood ratio test for branch support
MCMC: Markov chain Monte Carlo.

## Availability of supporting data

The datasets generated and/or analysed during the current study are available in the github repository, github.com/sebastianduchene/bacteria_genomic_rates_data

## Declarations

### Ethics and consent to participate

Not applicable.

### Consent to publish

Not applicable.

### Availability of data and materials

All the software used in this study is freely available and open source. The data are all available online github.com/sebastianduchene/bacteria_genomic_rates_data

### Author contributions

SD, DD, JLG, and KEH conceived and designed the experiments. SD, JLG, JH and ZD analysed the data. SD wrote the manuscript with input from all the authors.

## Acknowledgements

Not applicable.

## Supplementary material legends

**Fig.S1. Root-to-tip regression for all data sets.** The blue points correspond to tips in the tree. The black line represents the linear regression of root-to-tip distance as a function of the sampling time. The root-to-tip distance is measured by fitting the root of the tree that maximises R^2^.

## References

1. Gire SK, Goba A, Andersen KG, Sealfon RSG, Park DJ, Kanneh L, et al. Genomic surveillance elucidates Ebola virus origin and transmission during the 2014 outbreak. Science; 2014;345:1369–72.

2. Holmes EC, Dudas G, Rambaut A, Andersen KG. The evolution of Ebola virus: Insights from the 2013–2016 epidemic. Nature. 2016;538:193–200.

3. Drummond AJ, Suchard MA, Xie D, Rambaut A. Bayesian phylogenetics with BEAUti and the BEAST 1.7. Mol Biol Evol. 2012;29:1969–73.

4. Bouckaert R, Heled J, Kühnert D, Vaughan T, Wu C-H, Xie D, et al. BEAST 2: a software platform for Bayesian evolutionary analysis. PLOS Comput Biol. 2014;10:e1003537.

5. Ho SYW, Duchêne S. Molecular-clock methods for estimating evolutionary rates and time scales. Mol Ecol. 2014;23:5947–75.

6. Volz EM, Koelle K, Bedford T. Viral phylodynamics. PLOS Comput Biol. 2013;9:e1002947.

7. du Plessis L, Stadler T. Getting to the root of epidemic spread with phylodynamic analysis of genomic data. Trends Microbiol. 2015;23:383–6.

8. Rieux A, Balloux F. Inferences from tip-calibrated phylogenies: a review and a practical guide. Mol Ecol. 2016;25:1911–24.

9. Stadler T, Kouyos R, von Wyl V, Yerly S, Böni J, Bürgisser P, et al. Estimating the basic reproductive number from viral sequence data. Mol Biol Evol. 2012;29:347–57.

10. Wong VK, Baker S, Pickard DJ, Parkhill J, Page AJ, Feasey NA, et al. Phylogeographical analysis of the dominant multidrug-resistant H58 clade of Salmonella Typhi identifies inter-and intracontinental transmission events. Nat Genet. 2015;47:632–9.

11. Njamkepo E, Fawal N, Tran-Dien A, Hawkey J, Strockbine N, Jenkins C, et al. Global phylogeography and evolutionary history of Shigella dysenteriae type 1. Nat Microbiol. 2016;1:16027.

12. Stamatakis A. RAxML version 8: a tool for phylogenetic analysis and post-analysis of large phylogenies. Bioinformatics. 2014;30:1312–3.

13. Nguyen L-T, Schmidt HA, von Haeseler A, Minh BQ. IQ-TREE: a fast and effective stochastic algorithm for estimating maximum-likelihood phylogenies. Mol Biol Evol. SMBE; 2015;32:268–74.

14. Guindon S, Dufayard J-F, Lefort V, Anisimova M, Hordijk W, Gascuel O. New algorithms and methods to estimate maximum likelihood phylogenies: assessing the performance of PhyML 3.0. Syst Biol. 2010;59:307–21.

15. Price MN, Dehal PS, Arkin AP. FastTree 2–approximately maximum-likelihood trees for large alignments. PLoS One. 2010;5:e9490.

16. To T-H, Jung M, Lycett S, Gascuel O. Fast dating using least-squares criteria and algorithms. Syst Biol. 2016;65:82–97.

17. Duchêne S, Geoghegan JL, Holmes EC, Ho SYW. Estimating evolutionary rates using time-structured data: a general comparison of phylogenetic methods. Bioinformatics. 2016;32:3375–9.

18. Kumar S, Hedges SB. Advances in time estimation methods for molecular data. Mol Biol Evol. 2016;

19. Volz EM, Frost SDW. Scalable relaxed clock phylogenetic dating. Virus Evol. 2017;3.

20. Sagulenko P, Puller V, Neher R. TreeTime: maximum likelihood phylodynamic analysis. bioRxiv. 2017;153494.

21. Stadler T. Mammalian phylogeny reveals recent diversification rate shifts. Proc Natl Acad Sci. 2011;108:6187–92.

22. Höhna S, Landis MJ, Heath TA, Boussau B, Lartillot N, Moore BR, et al. RevBayes: Bayesian phylogenetic inference using graphical models and an interactive model-specification language. Syst Biol. 2016;65:726–36.

23. Poon AFY. Phylodynamic inference with kernel ABC and its application to HIV epidemiology. Mol Biol Evol. 2015;32:2483–95.

24. Saulnier E, Alizon S, Gascuel O. Assessing the accuracy of Approximate Bayesian Computation approaches to infer epidemiological parameters from phylogenies. PLOS Comput Biol. 2017;13:e1005416.

25. Merker M, Blin C, Mona S, Duforet-Frebourg N, Lecher S, Willery E, et al. Evolutionary history and global spread of the Mycobacterium tuberculosis Beijing lineage. Nat Genet. 2015;47:242–9.

26. Devault AM, Golding GB, Waglechner N, Enk JM, Kuch M, Tien JH, et al. Second-pandemic strain of Vibrio cholerae from the Philadelphia cholera outbreak of 1849. N Engl J Med. 2014;370:334–40.

27. Baines SL, Holt KE, Schultz MB, Seemann T, Howden BO, Jensen SO, et al. Convergent adaptation in the dominant global hospital clone ST239 of methicillin-Resistant Staphylococcus aureus. MBio. 2015;6:e00080–15.

28. Drummond AJ, Ho SYW, Phillips MJ, Rambaut A. Relaxed phylogenetics and dating with confidence. PLOS Biol. 2006;4:699–710.

29. Ho SYW, Duchêne S, Duchêne D. Simulating and detecting autocorrelation of molecular evolutionary rates among lineages. Mol Ecol Resour. 2015;15.

30. Duchêne S, Duchêne DA, Di Giallonardo F, Eden J-S, Geoghegan JL, Holt KE, et al. Cross-validation to select Bayesian hierarchical models in phylogenetics. BMC Evol Biol. 2016;16.

31. Anisimova M, Gascuel O. Approximate likelihood-ratio test for branches: A fast, accurate, and powerful alternative. Syst Biol. 2006;55:539–52.

32. Ramsden C, Holmes EC, Charleston MA. Hantavirus evolution in relation to its rodent and insectivore hosts: no evidence for codivergence. Mol Biol Evol. 2009;26:143–53.

33. Duchêne S, Duchêne DA, Holmes EC, Ho SYW. The performance of the date-randomization test in phylogenetic analyses of time-structured virus data. Mol Biol Evol. 2015;32:1895–906.

34. Rambaut A, Lam TT, Carvalho LM, Pybus OG. Exploring the temporal structure of heterochronous sequences using TempEst (formerly Path-O-Gen). Virus Evol. 2016;2:vew007.

35. Duchêne S, Holt KE, Weill F-X, Le Hello S, Hawkey J, Edwards DJ, et al. Genome-scale rates of evolutionary change in bacteria. Microb Genomics. Microbiology Society; 2016;2:e000094.

36. Boskova V, Bonhoeffer S, Stadler T. Inference of epidemiological dynamics based on simulated phylogenies using birth-death and coalescent models. PLOS Comput Biol. 2014;10:e1003913.

37. Rosenberg NA, Nordborg M. Genealogical trees, coalescent theory and the analysis of genetic polymorphisms. Nat Rev Genet. 2002;3:380.

38. Boskova V, Stadler T, Magnus C. The influence of phylodynamic model specifications on parameter estimates of the Zika virus epidemic. Virus Evol. 2018;4:vex044.

39. Murray GGR, Wang F, Harrison EM, Paterson GK, Mather AE, Harris SR, et al. The effect of genetic structure on molecular dating and tests for temporal signal. Methods Ecol Evol. 2015;7:80–9.

40. Stadler T, Kühnert D, Bonhoeffer S, Drummond AJ. Birth–death skyline plot reveals temporal changes of epidemic spread in HIV and hepatitis C virus (HCV). Proc Natl Acad Sci. 2013;110:228–33.

41. Kühnert D, Stadler T, Vaughan TG, Drummond AJ. Phylodynamics with migration: A computational framework to quantify population structure from genomic data. Mol Biol Evol. 2016;33:2102–16.

42. Croucher NJ, Page AJ, Connor TR, Delaney AJ, Keane JA, Bentley SD, et al. Rapid phylogenetic analysis of large samples of recombinant bacterial whole genome sequences using Gubbins. Nucleic Acids Res. 2014;43:e15.

43. Rambaut A, Grass NC. Seq-Gen: an application for the Monte Carlo simulation of DNA sequence evolution along phylogenetic trees. Bioinformatics. 1997;13:235–8.

